# Regulation of Human Erythroferrone Expression

**DOI:** 10.64898/2026.07.02.735786

**Authors:** Gemma Moir-Meyer, Robert Sertori, Cavan Bennett, Martin Pal, Anne Pettikiriarachchi, Jim Hughes, Hal Drakesmith, James O. J. Davies, Damien J. Downes, Matthew E. Gosden, Mohsin Badat, Danielle Clucas, Christian Babbs, Ryo Kurita, Connie S. N. Li-Wai-Suen, Alexandra L. Garnham, Natalia Benetti, Megan Iminitoff, Tamara Cameron, Marnie Blewitt, Sant-Rayn Pasricha

**Affiliations:** Walter and Eliza Hall Institute of Medical Research, Parkville, Victoria, Australia; Department of Medical Biology, University of Melbourne, Parkville, Victoria, Australia; School of Dentistry and Medical Sciences, Faculty of Science and Health, Charles Sturt University, Wagga Wagga, NSW, Australia; MRC Weatherall Institute for Molecular Medicine, University of Oxford, Oxford, United Kingdom; Centre for Genomics and Child Health, Blizard Institute, Queen Mary University of London, UK; Department of Clinical Haematology, The Royal Melbourne Hospital and the Peter MacCallum Cancer Centre, Parkville, Victoria, Australia; Department of Research and Development, Central Blood Institute, Blood Service Headquarters, Japanese Red Cross Society, Tokyo, Japan; Melbourne School of Population and Global Health, The University of Melbourne, Parkville, Victoria, Australia

**Author notes:** Correspondence: Sant-Rayn Pasricha, Walter and Eliza Hall Institute of Medical Research, 1G Royal Pde, Parkville, VIC 3052, Australia. GMM and RS contributed equally to this study. Data Sharing Statement: For original data, please contact the corresponding author.

## Abstract

Erythroferrone (ERFE) is an erythroblast-secreted hormone that suppresses hepatic hepcidin expression to increase iron availability for erythropoiesis, ensuring recovery from anaemia. ERFE excess drives iron overload in disorders of ineffective erythropoiesis. Despite its pivotal role in systemic iron homeostasis and diseases of erythropoiesis, ERFE’s molecular regulation has remained undefined. Here, we applied a genomic approach to characterise the molecular mechanisms governing *ERFE* expression. Using the HUDEP-2 human erythroid progenitor model, integrative ATAC-seq, CUT&RUN and micro capture-C analysis we identified a stage-specific accessible chromatin region within the *ERFE* 3’ UTR that interacts with the promotor. We also identified enhancer-associated chromatin marks including H3K4me1 and H3K27ac in this region, and demonstrate that this *cis*-regulatory element is bound by key erythroid transcription factors KLF1, GATA1, TAL1 and STAT5. Functional dissection using CRISPR-Cas9-mediated deletion of the central 3’ UTR enhancer element led to marked reduction in *ERFE* mRNA expression, and we show a corresponding reduction in nascent mRNA, confirming a key role for this region in transcriptional regulation. We define the transcriptional regulatory mechanism by which maturing human erythroblasts activate ERFE, the endocrine signal that coordinates erythropoietic demand with systemic iron mobilisation.

## Introduction

Erythropoiesis represents the body’s largest demand for iron. The availability of iron for cellular uptake is controlled by the systemic iron regulatory hormone, hepcidin, which is in turn transcriptionally regulated by infection and inflammation, hypoxia, iron sensing and erythropoiesis.^1^ Erythroferrone (encoded by *ERFE*) acts physiologically to suppress hepcidin^2,3^ (by operating as a decoy ligand for bone morphogenic protein 6 - BMP6)^4^ during erythroid expansion to enable recovery from anaemia.^5,6^ Increased erythroferrone also drives pathological iron overload in patients with chronic ineffective erythropoiesis such as ß-thalassaemia,^7^ and inhibition of erythroferrone is an attractive option to mitigate this problem.^8^

An improved understanding of how the body regulates *ERFE* expression could inform therapeutic opportunities for patients with iron overload. However, the molecular regulation of human *ERFE* gene expression has not been extensively described, beyond initial reports indicating a role for signal transducer and activator of transcription 5 (STAT5).^2^

Erythroid gene expression is orchestrated by stage-specific transcription factor networks often binding to distal enhancers with long-range chromatin interactions. Thus, unbiased genomic approaches offer important advantages over promoter-centric reporter assays for defining physiologically relevant regulatory mechanisms. For example, the field’s understanding of globin gene regulation has been driven by integrated genetic, epigenomic, three-dimensional chromatin assays and genome-editing studies that have identified enhancers controlling globin switching and foetal haemoglobin expression.^9,10^ Accordingly, mapping chromatin accessibility, enhancer-associated histone marks, transcription factor occupancy and promoter–enhancer contacts across erythroid maturation provides a powerful strategy to identify the regulatory elements that control erythroid genes in their native genomic context.

We therefore leveraged these strategies to discover the transcriptional control of erythroferrone, the erythroid hormone that governs systemic iron mobilisation during stress erythropoiesis and ineffective erythropoiesis. Here, we identify a novel regulatory element that drives the transcriptional regulation of this critical gene.

## Results

### ERFE transcription and chromatin landscape in human umbilical cord blood-derived erythroid progenitor 2 (HUDEP-2) cells

While mouse *Erfe* expression peaks in polychromatic erythroblasts^2^, *ERFE*’s transcriptional profile in human erythroblasts remains unknown. To quantify human *ERFE* mRNA abundance during erythropoiesis, quantitative PCR (qPCR) was performed on four erythroid cell stages isolated from HUDEP-2 cells differentiated *in vitro* and selected using fluorescence activated cell sorting (FACS) (Figure 1A-C). qPCR revealed that human *ERFE* expression is low in proerythroblasts, increasing in basophilic and maximal in polychromatic cells before falling in orthochromatic erythroblasts (Figure 1D), consistent with observations in mice.

**Figure 1:**
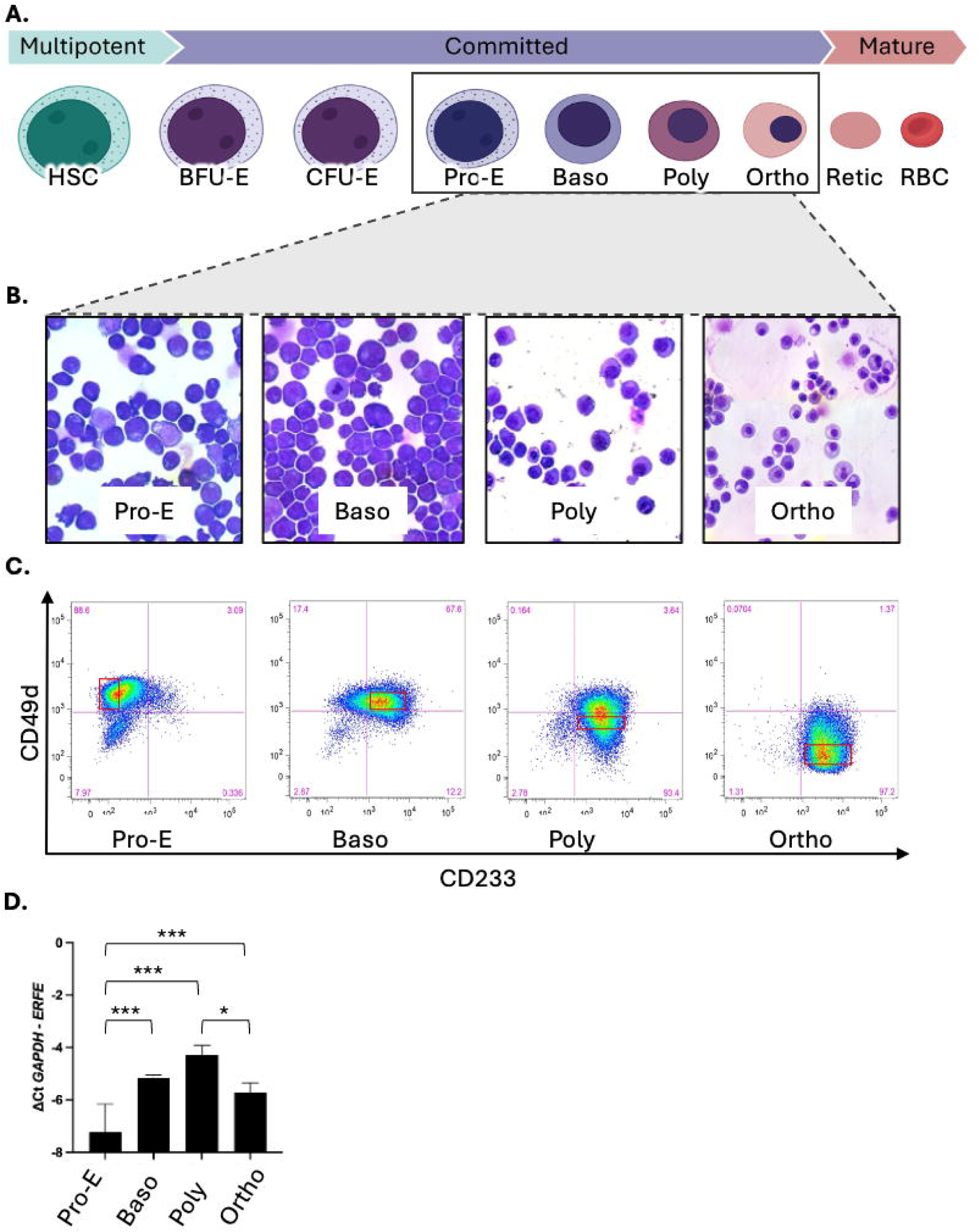
Human *ERFE* gene expression in different stages of HUDEP-2 *in vitro* erythropoiesis. HSC – Haematopoietic stem cell; BFU-E – burst forming unit erythroid; CFU-E – colony forming unit erythroid; Pro-E – proerythroblast; Baso – basophilic erythroblast; Poly – polychromatic erythroblast; Ortho – orthochromatic erythroblast. Late ortho – late orthochromatic erythroblast. (A) Black box indicates portion of erythropoiesis captured by the HUDEP-2 model system: definitive terminal erythropoiesis. (B) Cytospin images of flow cytometry-sorted HUDEP-2 erythroblast populations. (C) Flow cytometry sorting strategy used to isolate discrete populations of *in vitro* differentiated HUDEP-2 erythroblasts. Red squares denote gates for cells collected for downstream analysis. Panel shows expression of CD49d and CD233 on cells that are positive for CD235a. Pro-E (CD49d+/CD233-), Baso (CD49d+/CD233+), Poly (CD49d low/CD233+), and Ortho (CD49d-/CD233+). (D) Mean ±SEM difference in human *ERFE* expression calculated by ΔCt method relative to *GAPDH.* All qPCR assays were conducted with technical triplicates and biological replicates of n=5 or n=6. Statistical differences between groups were tested by repeated measure one way ANOVA and represented as: *\*\*\* P<0.001, * = P<0.05*

Next, we sought to evaluate the local chromatin landscape when *ERFE* expression is lowest (proerythroblasts) and highest (polychromatic erythroblasts). We performed assay for transposase-accessible chromatin (ATAC-seq) where DNA is treated with a hyperactive transposase that simultaneously cleaves and tags (with sequencing adaptors) open chromatin. When the same four distinct cell stages (Figure 2A) were interrogated, *ERFE*’s promoter was accessible from early erythropoiesis in proerythroblasts. Unexpectedly, we also observed high levels of chromatin accessibility (up to four distinct peaks) in *ERFE*’s 3’ untranslated region (UTR) and the downstream intergenic region near the adjacent gene ILK associated serine/threonine phosphatase (*ILKAP)*. Our data suggest the chromatin state changes over the *ERFE* 3’ region as erythroblasts mature, whereby proerythroblasts contain a single highly accessible region (peak 2), basophilic and polychromatic erythroblasts have an additional region of open chromatin (peaks 2 and 3), and orthochromatic erythroblasts have four punctate regions of open chromatin (peaks 1-4).

**Figure 2:**
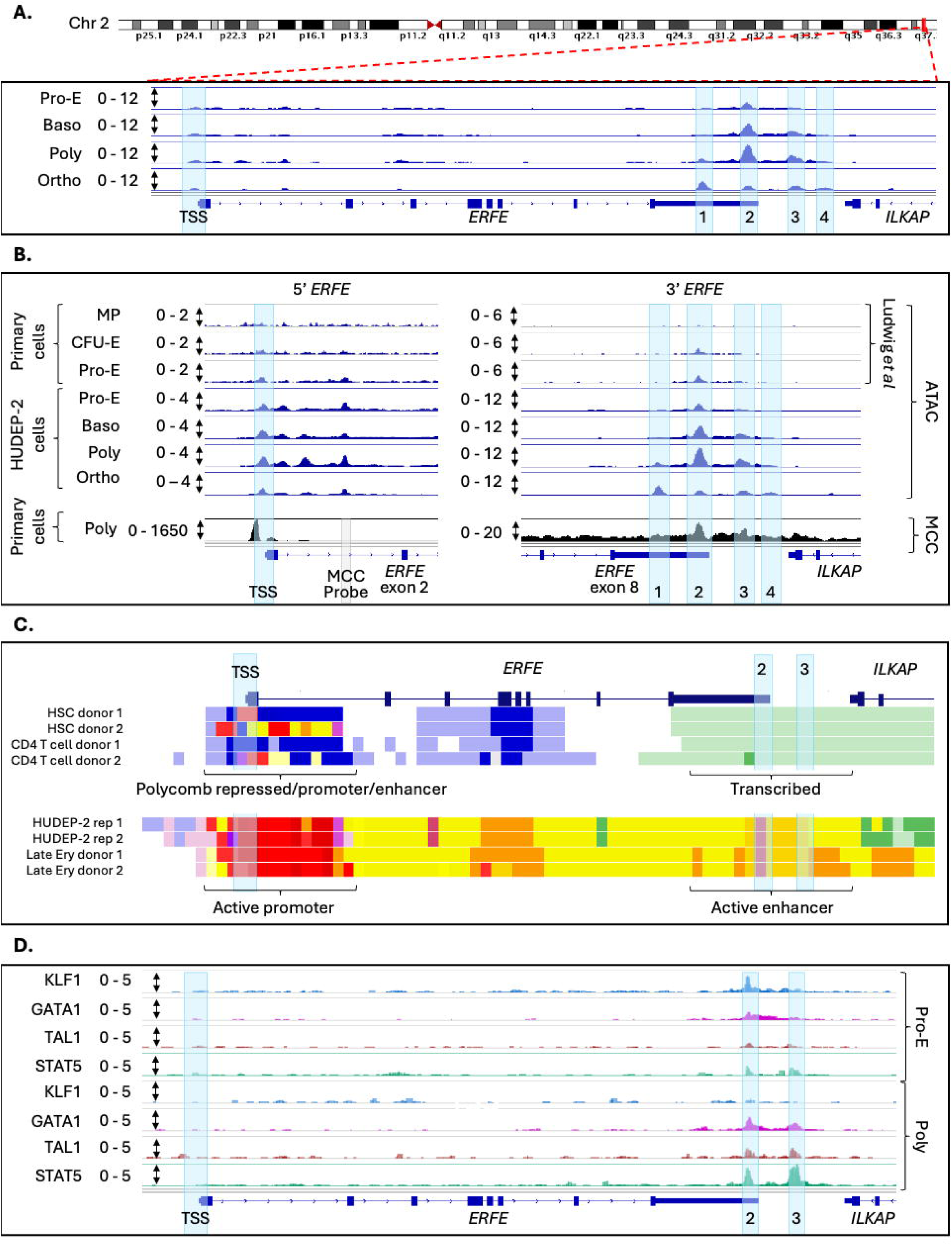
Enhancer-like architecture of the *ERFE* 3′ UTR with occupancy by erythroid transcription factors. Sequencing from proerythroblasts (pro-E), basophilic erythroblasts (baso), polychromatic erythroblasts (poly) and orthochromatic erythroblasts (ortho) where read depth (after spike in calibration for CUT&RUN) was normalised to counts per million reads and shown as a range on the y-axis. Scale was determined by the highest peak within the window across all four cell stages. Tracks are shown as an overlay of 2-3 replicates. Transcriptional start site (TSS) and enhancer peaks are shown with blue highlights. (**A**) *ERFE* locus highlighted in red on chromosome 2, and expanded to show ATAC-seq across four erythroid stages. (**B**) *ERFE’s* 5’ transcriptional start site and 3’ UTR are expanded to show timing of accessibility during erythroid maturation with data from Ludwig *et al.,* (2019). Micro Capture C (MCC) shows increased instances of contact between *ERFE*’s 5’ region and peaks 2 and 3 relative to the surrounding architecture in 3 biological replicates in erythroblasts differentiated from CD34+ cells from a single health donor. (**C**) UCSC Genome Browser image of data from Xiang *et al.,* (2024) showing epigenetic states at *ERFE* locus in haematopoietic stem cells (HSCs), CD4 T cells, HUDEP-2 erythroblasts and mature primary cell erythroblasts from *in vitro* differentiated CD34+ cells. Colours denote various epigenetic states derived by integrative modelling of ATAC-seq, CTCF occupancy, and levels of the histone modifications H3K27ac, H3K27me3, H3K36me3, H3K4me1, H3K4me3, and H3K9me3 (Xiang *et al*., 2024 – Fig 2B) in the UCSC Genome Browser track description. (**D**) CUT&RUN of KLF1, GATA1, TAL1 and STAT5 at *ERFE* locus in proerythroblasts and polychromatic erythroblasts (timepoints of minimal and maximal *ERFE* gene transcription).

Since *ERFE’s* promoter was already accessible in proerythroblasts and the HUDEP-2 model does not capture earlier haematopoietic lineage stages, we queried publicly accessible ATAC-seq generated from myeloid progenitor (MP) cells and colony forming units – erythroid (CFU-Es).^11^ These data showed that *ERFE*’s chromatin is closed in MPs and 3’ UTR peak-2 is the first region to become accessible at the CFU-E stage. Only after the CFU-Es mature to proerythroblasts do ATAC-seq peaks emerge at the promoter, suggesting that there may be regulatory elements in peak-2 whose activation precedes *ERFE*’s transcriptional initiation (Figure 2B).

We next sought to evaluate whether the accessible region within *ERFE*’s 3’UTR was associated with promoter accessibility, for instance via *cis*-activation. Using Micro Capture-C (MCC)^12^, a high-resolution chromatin capture technique that identifies DNA regions that are co-located in 3-dimensional space, we demonstrated an interaction between 5’ *ERFE* and 3’ UTR peaks 2 and 3 in polychromatic primary erythroblasts (Figure 2B). This indicates that a *cis*-element within the 3’ UTR interacts with the *ERFE* promoter region.

### Enhancer chromatin modifications overlap ERFE’s 3’ UTR

We next explored the regulatory events that coincide with, or precede, maximal *ERFE* transcription. Histone modifications are associated with chromatin compaction to either facilitate or inhibit the access of DNA binding proteins^13,14^ and we explored the *ERFE* locus in a large integrative modelling analysis.^15^ These data derived epigenetic states from the combination of ATAC-seq, CCCTC-binding factor (CTCF) binding and six histone modifications, and defined active erythroid promoters as being nuclease-accessible with moderate-high levels of histone H3 lysine 4 trimethylation (H3K4me3) and histone H3 lysine 27 acetylation (H3K27ac). Active erythroid enhancers were defined as nuclease-accessible with moderate-high levels of histone H3 lysine 4 monomethylation (H3K4me1) and H3K27ac. The 5’ *ERFE* region is annotated as an active promoter (validating this approach) and the 3’ UTR peaks 2 and 3 are annotated as an active enhancer region in erythroid and HUDEP-2 cells, but not in hematopoietic stem cells (HSC) or T cells, indicating tissue specificity of this putative enhancer region in erythroblasts (Figure 2C).

### ERFE’s 3’ UTR integrates erythroid and stress signalling

Next, we sought to define the transcription factor binding landscape in the 3’ UTR putative enhancer region. In peaks 2 and 3 we found transcription factor binding motifs for Krüppel-like factor 1 (KLF1), GATA-binding factor 1 (GATA1) and T-cell acute leukaemia protein 1 (TAL1). KLF1 is a pioneer transcription factor responsible for facilitating chromatin accessibility at major erythroid genes which can form complexes with GATA1 and TAL1 and divert these binding partners to erythroid activation sites.^16–18^ We also sought to understand whether there was colocalization of STAT5, a transcription factor mediating trans-activation of erythropoietin (via Janus kinase (JAK)-STAT) to facilitate stress erythropoiesis.

Based on this, we sought to explore the binding of these transcription factors at the *ERFE* region, in proerythroblasts and polychromatic erythroblasts. Cleavage under targets and release using nuclease (CUT&RUN) utilises a fusion protein comprising Protein A IgG, Protein G IgG and Micrococcal Nuclease (pAG-MNase) to bind a primary antibody and isolate DNA-protein hybrids that can be quantified with sequencing. CUT&RUN demonstrated differential binding of KLF1, GATA1, TAL1 and STAT5 to *ERFE*’s 3’ UTR. KLF1 was bound at peak-2 in proerythroblasts but not in polychromatic erythroblasts, while GATA1, TAL1 and STAT5 were bound at peaks 2 and 3, with increased abundance during maximal *ERFE* expression in polychromatic erythroblasts (Figure 2D). This may reflect chromatin remodelling facilitated by KLF1 in early *ERFE* induction, alongside subsequent signal integration with STAT5 to support maximal *ERFE* transcription.

### Clustered regularly interspaced short palindromic repeats (CRISPR) induced deletions in core binding motifs in the putative ERFE enhancer abrogate ERFE expression

Since peak-2 is *ERFE*’s first region to exhibit chromatin accessibility and transcription factor occupancy, we next aimed to assess its functional relevance in the gene’s activation and used genome editing to introduce insertion-deletions (indels) within transcription factor binding motifs. HUDEP-2 cells were transduced with CRISPR-associated protein 9 (Cas9) lentivirus and single guide RNAs (sgRNAs) targeting the peak-2 GATA1-TAL1 binding motif (JASPAR MA0140.3). Following clonal isolation and Sanger sequencing of the targeting site, two HUDEP-2 lines were isolated: ‘ERFE 1’ containing a homozygous 66 bp deletion removing KLF1 (JASPAR MA0493.3) and GATA1-TAL1 binding motifs and partial removal of STAT5 (JASPAR MA0519.1), and ‘ERFE 2, containing a homozygous 25 bp deletion which removed only the GATA1 and KLF1 motifs (Figure 3A).

**Figure 3:**
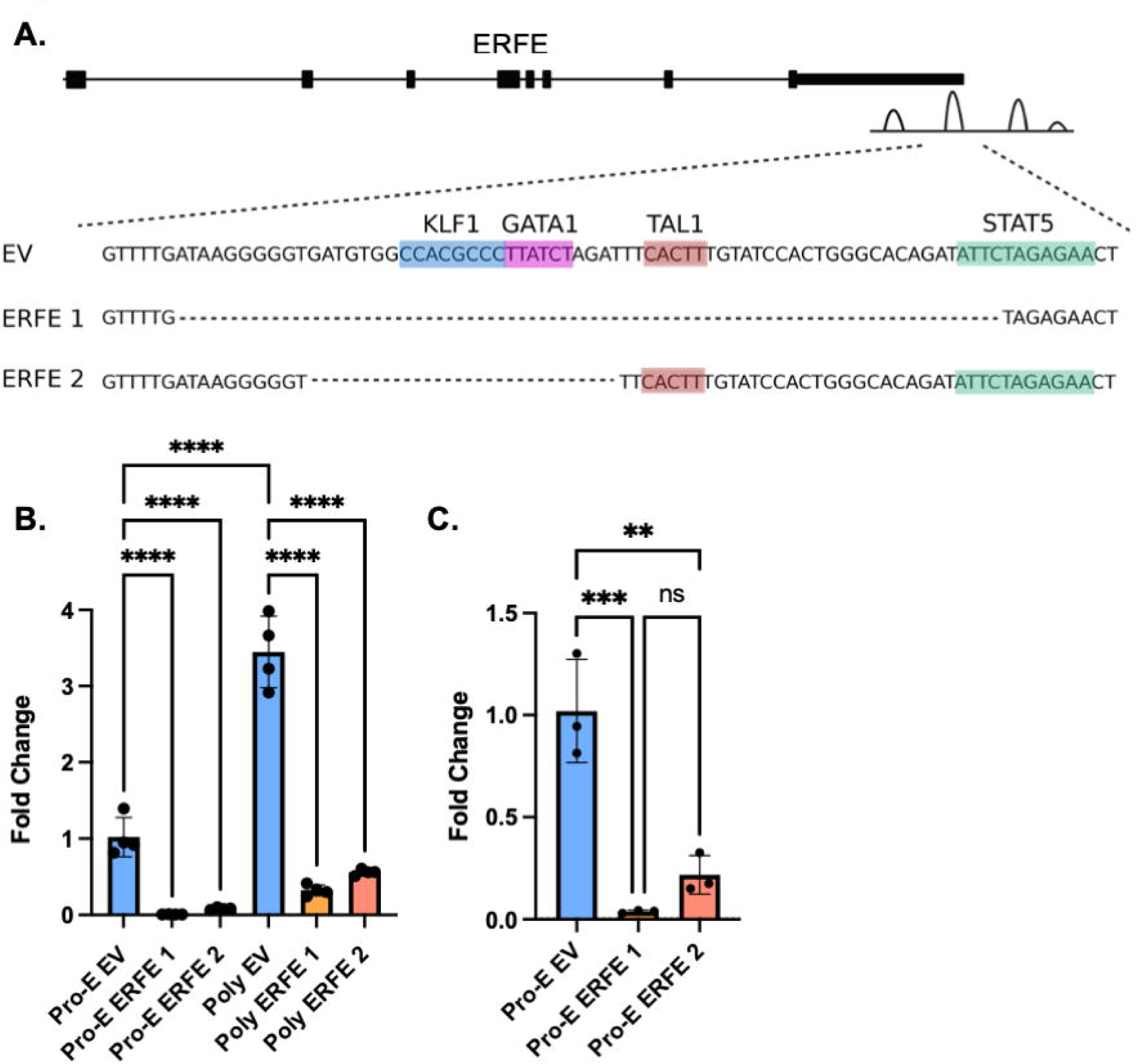
Biallelic disruption of transcription factor motifs in the putative *ERFE* enhancer abrogates *ERFE* mRNA expression. **(A)** Schematic representation of CRISPR-induced deletion of KLF1, GATA1, TAL1 and STAT5 binding sites identified by CUT&RUN as per Figure 2. Deletion indicated by dotted lines based on Sanger sequencing. (**B**) *ERFE* expression at pro- and polychromatic erythroblast stage with EV coloured blue, ERFE 1 orange and ERFE 2 pink. (**C**) Nascent *ERFE* expression at proerythroblast stage (EV coloured blue, ERFE 1 orange and ERFE 2 pink). Statistical differences between groups were tested by one way ANOVA and are represented as follows: ∗∗∗∗ = *P* < .0001; ∗∗∗ = *P* < .001; ∗∗ = *P* < .01 and ns, not significant.

*ERFE* mRNA expression was compared by qPCR between the deletion lines and a non-targeting sgRNA control line (empty vector - EV). At the proerythroblast stage, we observed a 113-fold (ERFE 1) and 12-fold (ERFE 2) reduction in *ERFE* mRNA expression relative to the EV control (Figure 3B). At the polychromatic erythroblast stage when *ERFE* expression is maximal, we observed an 11-fold and 6-fold respective reduction in expression (Figure 3B).

To confirm that this reduction in *ERFE* expression was due to reduced mRNA synthesis rather than altered mRNA transcript stability, we measured nascent RNA expression. RNA from proerythroblasts was labelled by 5-ethylene uridine and captured using the Click-iT Nascent RNA Capture Kit. We observed substantial reductions in nascent *ERFE* mRNA expression for ERFE 1 and ERFE 2 compared with the EV control (P < 0.001) (Figure 3C) indicating that reduction in *ERFE* expression is likely driven by decreased transcription, supporting the conclusion that peak-2 contains an enhancer region regulating *ERFE* expression.

## Discussion

Here, we define the first transcriptional regulatory architecture of *ERFE*, the gene encoding erythroferrone, the central erythroid regulator of systemic iron metabolism that drives iron overload in disorders of dyserythropoiesis including thalassaemia.^19^ By integrating chromatin accessibility, erythroid transcription factor occupancy, chromatin conformation and functional validation using CRISPR-Cas9 genome editing, we identify a *cis*-regulatory element within the 3′ untranslated region of *ERFE* that is required for its activation during erythropoiesis. These findings provide a mechanistic explanation for how maturing erythroblasts induce *ERFE* expression and thereby link erythropoietic activity to systemic iron mobilisation through hepcidin suppression. More broadly, our data establish *ERFE* regulation as part of convergent signalling programs, canonical and stress erythropoiesis, and provide a foundation for understanding how dysregulated erythroid signalling contributes to iron loading in disorders of ineffective erythropoiesis.

To date, understanding of *ERFE* transcriptional regulation has been limited. The initial description of erythroferrone established that *ERFE/Fam132b* is rapidly induced in the erythroblast compartment of the bone marrow in mice, in response to erythropoietic stimulation, with pharmacological pathway-inhibition experiments implicating EPO–JAK2–STAT5 signalling in this response.^2^ Initial promotor analysis identified a putative STAT5 response element upstream of the gene,^2^ but subsequent work to identify the regulation of *ERFE in vivo* or *in vitro* has been limited. Functionally, ERFE acts as an erythroblast-derived endocrine signal that suppresses hepatic hepcidin production, thereby increasing intestinal iron absorption and mobilising iron from stores to support haemoglobin synthesis during stress erythropoiesis. Subsequent work has shown that ERFE suppresses hepcidin by antagonising hepatic BMP– Suppressor of Mothers Against Decapentaplegic (SMAD) signalling.^4,8^ However, despite this central role in iron homeostasis, the transcription factor networks that control *ERFE* activation have remained undefined.

*ERFE’*s activation is particularly pronounced following blood loss or haemolysis to enable recovery from anaemia during a process of rapid erythron expansion termed stress erythropoiesis.^20^ EPO is the primary regulator of red cell synthesis. When EPO binds to EPO-receptor (EPOR) on CFU-Es it activates proliferation and cell-survival pathways and signals the erythroid transcription program involving KLF1, GATA1 and TAL1.^21^ Despite EPO levels increasing 1000-fold during hypoxic stress,^20^ EPO concentration alone cannot trigger the stress erythropoiesis program which requires STAT5 phosphorylation to instead modify EPO responsiveness.^22^ These convergent signalling pathways are reflected in colocation of STAT5 with key erythroid transcription factors in the *ERFE* 3’UTR cis-regulatory element. These findings are consistent with previously observed patterns of colocalization of STAT-5 with erythroid master regulators at enhancer elements.^23^

The identification of a regulatory element within the 3′ UTR of *ERFE* is consistent with the increasingly recognised importance of *cis*-regulatory architecture in erythroid gene control. This region acquired progressive chromatin accessibility during erythroid maturation, physically interacted with the *ERFE* promoter, and was occupied by GATA1, KLF1, TAL1 and STAT5 in a stage-specific manner. In contrast to mouse data which reported a STAT5 binding motif upstream of *ERFE*’s promoter, our human data does not support 5’ binding of STAT5 and instead demonstrates its involvement in the 3’ regulatory region. The marked reduction in *ERFE* expression following CRISPR-Cas9 disruption of this putative enhancer region provides functional evidence that peak-2 is required for appropriate *ERFE* activation during human erythropoiesis. This mechanism is consistent with the broader paradigm of enhancer–promoter cooperation in fine-tuning gene expression during erythropoiesis.^24^

The 3′ untranslated region is classically recognised as a post-transcriptional regulatory region, containing sequence elements that influence mRNA stability, localisation, polyadenylation and translational efficiency. It also frequently harbours binding sites for microRNAs and RNA-binding proteins, enabling dynamic control of transcript abundance and protein output.^25^ In the context of *ERFE* regulation, our data suggest that the *ERFE* 3′ UTR has an additional DNA-level regulatory function: containing an enhancer-like element that acquires chromatin accessibility during erythroid maturation, is occupied by EPO-responsive transcription factors, and forms a local chromatin interaction with the *ERFE* promoter.

Collectively, our data support a model in which *ERFE* induction is not driven solely by promoter activation, but by engagement of a distal enhancer-like element embedded within the 3′ untranslated region.

## Acknowledgments

GMM was funded by a Melbourne Research Scholarship. SRP is funded by National Health and Medical Research Council of Australia (NHMRC) Fellowship GNT2009047. MEB was funded by NHMRC Investigator grants 1194345 and 2041117. MI and NB were funded by Australian Research Training Program Scholarships. The contents of this work are the responsibility of the authors and do not reflect the views of the NHMRC. This work was also supported by the Victoria State Government’s Operational Infrastructure Support Program and the NHMRC Independent Research Institute Infrastructure Support Scheme (IRIISS).

## Authorship Contributions

GMM and SP conceived the study. GMM, RS, CB, MP, AP, DJD, MEG, MB, DC, CBa undertook experiments, and GMM, RS, JRH, JD, MEG, CSNLWS, AG, NB, MI and TC analysed data. GMM, RS, JRH, HD, JOJD, MB and SP interpreted the data. RK contributed critical materials to the study. GMM, RS and SP drafted the manuscript, and all authors approved the final manuscript.

## Methods

### Cell Culture

HUDEP-2 cells were obtained from RIKEN and cultured as previously reported^26^, differentiated *in vitro* and collected at proerythroblast, basophilic, polychromatic and orthochromatic stages. Differentiation was validated by cell morphology using May-Grünwald and Giemsa staining, and immunophenotyping for CD49d (Abcam ab91051-PE), CD71 (Becton Dickinson Biosciences 563671-APC), CD233 (National Health Service Blood and Transplant International Blood Group Reference Laboratory, NHSBT IBGRL, 9439-FITC), CD235a (Abcam ab91163-APC), and propidium iodide (Abcam ab14083). Primary CD34+ cells were isolated from a healthy adult donor and differentiated as previously described.^27^

### qPCR

qPCR was used to measure the expression of various erythroid genes during maturation. RNA was isolated from HUDEP-2 cells using ISOLATE II RNA Kit and cDNA was prepared using SensiFAST cDNA Synthesis Kit (both Meridian Bioscience) following manufacturer’s instructions. (qPCR was performed in triplicate (LightCycler 480) using TaqMan assays and SensiFAST Probe No Rox Kit (Meridian Bioscience). Expression was normalised to *GAPDH* and calculated using the ΔCT method for Figure 1, and ΔΔCT method for Figure 3, using Taqman probes (Thermo Fisher Scientific) ERFE Hs00937727_m1, GAPDH Hs04420632_g1 and Hs02758991_g1.

### ATAC-Sequencing

Chromatin accessibility was assayed genome-wide using ATAC-sequencing. Sorted HUDEP-2 cells (7.5 × 10⁴) were collected by flow cytometry, pelleted and washed, and processed using.^28^ Libraries were purified by Qiagen MinElute pre-PCR, AMPure XP post-PCR (Beckman Coulter), quantified and quality-checked on the Agilent Tapestation. Pooled libraries were sequenced (150bp paired end) on the Illumina NextSeq. Reads were quality-checked (FastQC, MultiQC), trimmed (Trim Galore), and aligned to hg38 (Rsubread). BAM files were sorted and deduplicated (Sambamba), and read depth was normalised to counts per million before files were converted to bigWig format.

### Micro Capture-C (MCC)

Micro Capture-C enables 3-dimensional chromatin architecture at loci of interest to be probed at base pair resolution. Formaldehyde crosslinking fixes the DNA in its native 3D state to capture chromatin regions in proximity. Following cell permeabilization, MNase treatment cleaves the DNA in a sequence-independent manner, and the fragments undergo library preparation after protein digestion, decrosslinking and purification. MCC libraries from 1 x 10^7^ day 10 CD34+ cells were generated, sequenced and analysed as described.^29^ *ERFE* promotor interactions were probed using the following sequence: GGGAGATCAGCCAGCAGGTGGTGCAGTTGACCCTCTAAGGCACAAGGAAGCACTGAGCAGCGCTGAGGGCCCTGGGCCCCTGAACCCTTTCTCCGCCCAACCACAGGCTCCTCGGCACCG

### CUT&RUN

CUT&RUN was used to assay protein binding to genomic elements of interest. Unlike chromatin immunoprecipitation (ChIP), CUT&RUN can be performed on freshly isolated nuclei without crosslinking or chromatin shearing. Following selection by flow cytometry, 5 × 10⁵ cells were processed with the CUTANA Kit (EpiCypher) according to the manufacturer’s protocol using the following antibodies: Abcam GATA1 (ab11852), TAL1 (ab155195); Atlas KLF1 (HPA-051850), Thermo Fisher Scientific STAT5A (13-3600), IgG-antibody control Epicypher (13-0042k), and a no-antibody control. Sequencing and data processing was performed as for ATAC-seq.

### CRISPR-Cas9

Functional validation of the genomic elements of interest was assayed through targeted deletions. HUDEP-2 cells expressing Cas9-GFP were transduced with either an empty vector control (sgRNA.SFFV.tBFP; Addgene #169940) or sgRNAs targeting the GATA1–TAL1 binding motif in peak 2 5′-GATTTCACTTTGTATCCACT-3′, and 5′-GTGAAATCTAGATAAGGGCG-3′). Clonal lines were established by single-cell sorting of GFP⁺/BFP⁺ cells, and editing was confirmed by Sanger sequencing.

### Click-iT Nascent RNA

To assay newly transcribed (nascent) mRNA transcription, HUDEP-2 cells were labelled with 0.2mM 5-ethynyl uridine for 2 hours. RNA was extracted according to Isolate II RNA mini kit (Meridian Bioscience). 1 μg total RNA was biotinylated with 0.5mM biotin azide and precipitated with glycogen, ammonium acetate and ethanol. RNA concentration measured by Ds11 spectrometer (DeNovix). 400ng of biotinylated RNA was bound to 25ul Dynabeads MyOne Streptavidin T1 magnetic beads and washed according to the protocol. Bead were resuspended in 15µl of H2O. Bead/RNA mixture was added to SensiFAST cDNA synthesis mix (Meridian Bioscience), and the transcribed cDNA used for qPCR.

## References

1 Camaschella, C., Pagani, A., Silvestri, L. & Nai, A. The mutual crosstalk between iron and erythropoiesis. Int J Hematol 116, 182–191 (2022). 10.1007/s12185-022-03384-y

2 Kautz, L. et al. Identification of erythroferrone as an erythroid regulator of iron metabolism. Nat Genet 46, 678–684 (2014). 10.1038/ng.2996

3 Coffey, R. et al. Erythroid overproduction of erythroferrone causes iron overload and developmental abnormalities in mice. Blood 139, 439–451 (2022). 10.1182/blood.2021014054

4 Arezes, J. et al. Erythroferrone inhibits the induction of hepcidin by BMP6. Blood 132, 1473–1477 (2018). 10.1182/blood-2018-06-857995

5 Pasricha, S. R., McHugh, K. & Drakesmith, H. Regulation of Hepcidin by Erythropoiesis: The Story So Far. Annu Rev Nutr 36, 417–434 (2016). 10.1146/annurev-nutr-071715-050731

6 Kautz, L., Jung, G., Nemeth, E. & Ganz, T. Erythroferrone contributes to recovery from anemia of inflammation. Blood 124, 2569–2574 (2014). 10.1182/blood-2014-06-584607

7 Kautz, L. et al. Erythroferrone contributes to hepcidin suppression and iron overload in a mouse model of β-thalassemia. Blood 126, 2031–2037 (2015). 10.1182/blood-2015-07-658419

8 Arezes, J. et al. Antibodies against the erythroferrone N-terminal domain prevent hepcidin suppression and ameliorate murine thalassemia. Blood 135, 547–557 (2020). 10.1182/blood.2019003140

9 Deng, W. et al. Reactivation of developmentally silenced globin genes by forced chromatin looping. Cell 158, 849–860 (2014). 10.1016/j.cell.2014.05.050

10 Bauer, D. E. et al. An erythroid enhancer of BCL11A subject to genetic variation determines fetal hemoglobin level. Science 342, 253–257 (2013). 10.1126/science.1242088

11 Ludwig, L. S. et al. Transcriptional States and Chromatin Accessibility Underlying Human Erythropoiesis. Cell Rep 27, 3228–3240.e3227 (2019). 10.1016/j.celrep.2019.05.046

12 Hua, P. et al. Defining genome architecture at base-pair resolution. Nature 595, 125–129 (2021). 10.1038/s41586-021-03639-4

13 Kang, Y., Kim, Y. W., Kang, J. & Kim, A. Histone H3K4me1 and H3K27ac play roles in nucleosome eviction and eRNA transcription, respectively, at enhancers. Faseb j 35, e21781 (2021). 10.1096/fj.202100488R

14 Igolkina, A. A. et al. H3K4me3, H3K9ac, H3K27ac, H3K27me3 and H3K9me3 Histone Tags Suggest Distinct Regulatory Evolution of Open and Condensed Chromatin Landmarks. Cells 8 (2019). 10.3390/cells8091034

15 Xiang, G. et al. Interspecies regulatory landscapes and elements revealed by novel joint systematic integration of human and mouse blood cell epigenomes. Genome Res 34, 1089–1105 (2024). 10.1101/gr.277950.123

16 Wontakal, S. N. et al. A core erythroid transcriptional network is repressed by a master regulator of myelo-lymphoid differentiation. Proc Natl Acad Sci USA 109, 3832–3837 (2012). 10.1073/pnas.1121019109

17 Perkins, A. et al. Kruppeling erythropoiesis: an unexpected broad spectrum of human red blood cell disorders due to KLF1 variants. Blood 127, 1856–1862 (2016). 10.1182/blood-2016-01-694331

18 Magor, G. W. et al. KLF1 Acts As a Pioneer Transcription Factor Via SMARCA4 to Open Chromatin and Facilitate Redeployment of an Enhancer Complex Containing GATA1 and SCL. Blood 140, 696–697 (2022). 10.1182/blood-2022-157901

19 Kautz, L. et al. Erythroferrone contributes to hepcidin suppression and iron overload in a mouse model of beta-thalassemia. Blood 126, 2031–2037 (2015). 10.1182/blood-2015-07-658419

20 Socolovsky, M. Molecular insights into stress erythropoiesis. Curr Opin Hematol 14, 215–224 (2007). 10.1097/MOH.0b013e3280de2bf1

21 Cantor, A. B. & Orkin, S. H. Transcriptional regulation of erythropoiesis: an affair involving multiple partners. Oncogene 21, 3368–3376 (2002). 10.1038/sj.onc.1205326

22 Porpiglia, E., Hidalgo, D., Koulnis, M., Tzafriri, A. R. & Socolovsky, M. Stat5 signaling specifies basal versus stress erythropoietic responses through distinct binary and graded dynamic modalities. PLoS Biol 10, e1001383 (2012). 10.1371/journal.pbio.1001383

23 Gillinder, K. R. et al. Direct targets of pSTAT5 signalling in erythropoiesis. PLoS One 12, e0180922 (2017). 10.1371/journal.pone.0180922

24 Perreault, A. A. & Venters, B. J. Integrative view on how erythropoietin signaling controls transcription patterns in erythroid cells. Curr Opin Hematol 25, 189–195 (2018). 10.1097/moh.0000000000000415

25 Mayr, C. What Are 3’ UTRs Doing? Cold Spring Harb Perspect Biol 11 (2019). 10.1101/cshperspect.a034728

26 Kurita, R. et al. Establishment of immortalized human erythroid progenitor cell lines able to produce enucleated red blood cells. PLoS One 8, e59890 (2013). 10.1371/journal.pone.0059890

27 Scott, C. et al. Recapitulation of erythropoiesis in congenital dyserythropoietic anaemia type I (CDA-I) identifies defects in differentiation and nucleolar abnormalities. Haematologica 106, 2960–2970 (2021). 10.3324/haematol.2020.260158

28 Corces, M. R. et al. An improved ATAC-seq protocol reduces background and enables interrogation of frozen tissues. Nat Methods 14, 959–962 (2017). 10.1038/nmeth.4396

29 Hamley, J. C., Li, H., Denny, N., Downes, D. & Davies, J. O. J. Determining chromatin architecture with Micro Capture-C. Nat Protoc 18, 1687–1711 (2023). 10.1038/s41596-023-00817-8

